# Acute inflammation-mediated attenuation of behavioural sensitization in methamphetamine-sensitized mice via distinct COX-2 and TNF-α pathways

**DOI:** 10.64898/2026.05.08.723429

**Authors:** Rikuto Christopher Shinohara, Shuhei Ishikawa, Risa Matsumoto, Koki Ito, Mariko Tonosaki, Shiina Matsuyama, Masahiro Ohgidani, Minori Koga, Naoki Hashimoto, Ichiro Kusumi, Takahiro A. Kato

## Abstract

**Background and Purpose:** While inflammation has been generally considered to exacerbate symptoms of schizophrenia, some clinical observations suggest that acute inflammation may alleviate positive symptoms. However, animal models often use excessive inflammatory stimuli, and the effects of acute inflammation—comparable to levels observed in patients—remain unknown.

**Experimental Approach:** To address this, we examined whether acute inflammation induced under relatively mild, clinically relevant conditions suppresses behavioural sensitization in methamphetamine (METH)-sensitized mice, a model of psychostimulant-induced psychosis with relevance to certain aspects of positive symptoms of schizophrenia. We used a repeated METH (1 mg/kg) sensitized model to evaluate the effects of acute inflammation on behavioural sensitization. Acute inflammation was induced via two methods using either lipopolysaccharides (LPS; 1 μg/kg) to mimic peripheral immune activation or restraint stress (RS; single 2-h exposure) to model the neuroinflammation induced by psychological stress. LPS doses were adjusted with reference to the magnitude of peripheral cytokine elevation reported in patients, and RS was applied in short single sessions to avoid excessive inflammation.

**Key Results:** Both LPS and RS significantly suppressed behavioural sensitization, without inducing other behavioural abnormalities. This suppression was dependent on toll-like receptor-4 activation. LPS-mediated suppression involved cyclooxygenase-2, whereas RS-mediated suppression was linked to the microglia-derived tumour necrosis factor-α. LPS did not alter, whereas RS significantly reduced the striatal extracellular dopamine levels.

**Conclusion and Implications:** These findings suggest that acute inflammation suppresses behavioural sensitization through distinct mechanisms depending on the inflammatory trigger, providing a framework for understanding how inflammation may influence psychosis-related processes, with potential relevance to schizophrenia.

## Introduction

Schizophrenia is a psychiatric disorder characterized by positive symptoms such as hallucinations and delusions, which are the primary targets of pharmacological treatment.^(1)^ Neuroinflammation plays a role in the pathophysiology of schizophrenia, as patients with schizophrenia often exhibit elevated proinflammatory cytokine levels and microglial reactivity.^(2–6)^ Given these findings, inflammation has been considered a potential contributor to symptom exacerbation and a target for therapeutic intervention.^(7)^ However, some clinical studies have suggested that inflammation exerts protective effects under specific conditions.^(8–10)^ Historical evidence, such as the Wagner-Jauregg fever therapy, indicates that deliberate infection with malaria alleviates psychotic symptoms.^(8)^ Recent clinical studies have also reported temporary remission of positive symptoms following bacterial infection in patients with schizophrenia.^(9, 10)^ These findings suggest that acute inflammation does not always exacerbate schizophrenia symptoms and may even alleviate positive symptoms under specific circumstances.

In animal studies, psychostimulant-induced behavioural sensitization is widely employed as a model of psychostimulant-induced psychosis, where the observed sensitization serves as a surrogate marker for evaluating therapeutic agents, with relevance to certain aspects of the positive symptoms of schizophrenia.^(11–13)^ In studies using this model, peripheral acute inflammation induced by lipopolysaccharides (LPS), as well as other toll-like receptor 4 (TLR4) agonists, has been shown to suppress behavioural sensitization.^(14, 15)^ These findings may provide insights into the mechanisms underlying acute inflammation-associated symptom improvement in clinical settings. However, these studies have typically employed relatively high doses of LPS, and the resulting behavioural changes are often difficult to distinguish from sickness behaviour, potentially limiting their translational validity. Therefore, careful dose selection is required to induce a mild peripheral inflammatory response that approximates clinical conditions while minimally affecting baseline behaviour.

In addition to inducing peripheral immune activation with LPS, we also examined central neuroimmune responses as an alternative approach. Restraint stress (RS) is widely used in animal studies to induce stress-related neuroimmune responses via microglial reactivity and related pathways,^(16, 17)^ supporting its utility as a model of stress-induced neuroimmune activation. However, unlike LPS, the effects of RS-induced responses on behavioural sensitization remain unclear and warrant further investigation. Moreover, as the intensity and duration of stress critically influence these responses, RS was applied under conditions that avoid excessive stress.

Taken together, it remains unclear how acute inflammation of clinically relevant magnitude influences psychosis-related phenotypes. In the present study, we modelled mild acute inflammation based on clinical observations^(18)^, and examined its effects on methamphetamine (METH)-induced behavioural sensitization. To determine the specificity of these effects, we further assessed multiple behavioural domains. We aimed to clarify the impact of acute inflammation on behavioural sensitization, to elucidate the underlying mechanisms, and to test the hypothesis that inflammatory responses exert potentially protective effects.

## Materials and Methods

### Animals

Six-week-old male C57BL/6JJmsSlc mice (Japan SLC, Shizuoka, Japan) were housed under standard laboratory conditions (23 ± 1 °C; 50 ± 10% humidity; 12/12-h light/dark cycle; lights on at 8:00 AM, off at 8:00 PM), with free access to food and water. All experiments were approved by the Animal Research Committee (permission number: 21-0004) and adhered to the Guide for the Care and Use of Laboratory Animals of Hokkaido University (Sapporo, Japan).

### Drugs and treatments

METH (Dainippon Pharmaceutical, Tokyo, Japan) was dissolved in saline (Otsuka Pharmaceutical, Tokyo, Japan) and subcutaneously (s.c.) administered to the mice. Briefly, the mice were administered repeated METH injections (1 mg/kg/day) for five days (days 1–5). After a seven-day withdrawal period, the mice received a challenge dose of METH (1 mg/kg), and their locomotor activity was assessed. METH dose was determined based on a previous report.^(14)^ LPS (Merck KGaA, Darmstadt, Germany) was dissolved in saline, and intraperitoneally (i.p.) injected at a dose of 1 μg/kg 4 h prior to locomotor activity measurement. The LPS dose was determined based on preliminary experiments (Fig. S1A) and selected as the minimal dose that produced detectable peripheral immune activation, with reference to the magnitude of cytokine differences between healthy controls and patients with first-episode psychosis reported in clinical studies.^(18)^ Detailed methods are provided in Supplemental Methods. The interval between LPS administration and behavioural testing was set to 4 h to ensure sufficient elevation of the inflammatory cytokine levels, as previously described.^(18)^ As a TLR-4 inhibitor, TAK-242 (Merck KGaA, Darmstadt, Germany) was applied during the rescue experiment. TAK-242 (3 mg/kg) was diluted with phosphate-buffered saline (PBS) containing 0.9% dimethyl sulfoxide (FUJIFILM Wako Pure Chemical, Osaka, Japan) and administered i.p. 4 h prior to locomotor activity measurement.^(19)^ As a cyclooxygenase (COX)-2 inhibitor, nimesulide (Tokyo Chemical Industry, Tokyo, Japan) was applied in the rescue experiment. Nimesulide (10 mg/kg) was diluted with 0.2 mol/L Tris-HCl buffer solution (pH 8.0) and administered i.p. 4 h prior to locomotor activity measurement.^(20)^ As a TNF-α inhibitor, etanercept (Takeda Pharmaceutical, Tokyo, Japan), a TNF receptor 2-Fc fusion protein, was used in the rescue experiment. Etanercept (30 mg/kg) was reconstituted, diluted with Milli-Q water (Merck KGaA, Darmstadt, Germany), and administered s.c. on days 1, 3, and 5 prior to the rescue experiment, as previously described.^(21)^ All drugs were administered at a dose of 10 mL/kg.

### Restraint Stress

For stress experiments, the mice were restrained in 50-mL conical centrifuge tubes modified with small air holes to allow adequate ventilation for 2 h at 9:00 a.m., followed by a 2-h recovery period prior to behavioural tests. The detention time was set based on a preliminary report.^(22)^ The timing of RS initiation was set to 4 h before behavioural testing to match that of LPS administration.

### Behavioural tests

All behavioural tests were performed during daytime. Saline or LPS was administered or 2-h RS exposure was initiated 4 h before each behavioural test. Most behavioural tests were conducted in separate cohorts of mice to avoid potential carry-over effects between assays, although locomotor activity and the forced swim test were performed in the same mice to evaluate baseline activity.

### Locomotor activity

Home cage of each mouse was moved and placed under the sensor. Locomotor activity was measured using an infrared sensor detecting thermal radiation in mice using Supermex (Muromachi Kikai, Tokyo, Japan). Horizontal movements of the mice were digitized and fed to a computer every 5 min. The locomotor activity measurement apparatus was validated in our previous studies.^(23, 24)^ METH-induced locomotor activity was measured for 3 h following the administration of METH at 1 mg/kg after 2 h of environmental habituation. Spontaneous locomotor activity was measured for 10 min without environmental habituation under the same settings. Spontaneous locomotor activity was evaluated as the baseline motor ability.

### Forced swim test

Next, forced swim test was performed as previously described,^(25)^ with slight modifications. Briefly, each mouse was placed in a transparent glass cylinder (height: 18 cm; diameter: 11 cm) filled with water at 22–23 °C to a depth of 13 cm and forced to swim for 6 min. Immobility duration was measured using the SMART 3.0 video tracking software (Panlab, Barcelona, Spain). As indicated in a previous report,^(26)^ the forced swim test was used as a surrogate marker to evaluate the negative symptom-like behaviour of schizophrenia or depressive-like behaviour.

### Y-maze test

Y-maze test was conducted as previously described,^(21)^ with minor modifications. The Y-maze consisted of three arms, each being 40 cm long, 3 cm wide, and 12 cm deep, diverging 120° from a central point. Briefly, each mouse was placed in one arm and allowed to explore freely for 8 min. The maze was cleaned with 70% ethanol before each trial. The number of entries and sequence of arm entries were recorded. Additionally, number of alternations (successive entries into different arms) was divided by the total number of possible alternations (number of arms entered – 2) and multiplied by 100.^(27)^ The Y-maze test was used as a surrogate marker to assess cognitive function.^(28)^

### Surgery and microdialysis

The mice were anesthetized using a mixture of medetomidine (0.75 mg/kg, i.p.), midazolam (4 mg/kg, i.p.), and butorphanol (5 mg/kg, i.p.) and placed in a stereotaxic apparatus. Under anesthesia, G-4 guide cannula (Eicom, Kyoto, Japan) was stereotaxically implanted to target the surface of the striatum (coordinates: 1.0 mm anterior and 2.0 mm lateral to bregma and 3.3 mm below pia), as previously described.^(29)^ Then, FX-I-4-01 (Eicom) dialysis probe was inserted into the guide cannula, with 3.0 mm of the probe exposed to the striatal tissue. Microdialysis was performed on freely moving mice 10–14 d after surgery. Artificial cerebrospinal fluid (145 mM NaCl, 3.0 mM KCl, 1.3 mM CaCl_2_, and 1.0 mM MgCl_2_) was used, and perfusion was started 5 h before METH administration at a flow rate of 2 μL/min. After an initial 1-h perfusion, dialysis samples were collected every 20 min for 6 h into vials containing 50 μL of 50 mM acetic acid.

### Chromatographic analysis of the brain microdialysates

Extracellular dopamine levels were measured as previously described.^(24)^ HTEC-500 (Eicom) high-performance liquid chromatography system consisting of a reverse-phase ODS column with Eicompak PP ODS 30 4.6 mm was used to measure the dopamine levels. Dopamine levels were analyzed by injecting 20 μL of dialysate into the high-performance liquid chromatography system. Mobile phase consisted of 0.1 M phosphate buffer (PB; pH 6.0), 1% methanol (v/v), 50 mg/L Na_2_ ethylenediaminetetraacetic acid, and 500 mg/L sodium l-decanesulfonate. Separation was done at 25 °C at a flow rate of 0.5 mL/min. The electrochemical detector was set at an oxidation potential of 550 mV. Dopamine concentrations in the samples were calculated by comparing the areas with the standard dopamine solution. All data were analyzed using a data processor (EPC-500; Eicom) and the PowerChrom software program (eDAQ Pty, Sydney, Australia).

### Reverse transcription-quantitative polymerase chain reaction (RT-qPCR)

Mice were euthanized by cervical dislocation at the indicated time points following RS exposure, and the striatum was rapidly dissected. Cervical dislocation was performed exclusively by personnel who were appropriately trained and experienced, ensuring rapid and effective euthanasia. Every effort was made to minimize animal suffering. Total RNA was extracted from the striatum using the FastGene RNA Basic kit (NIPPON genetics, Tokyo, Japan), reverse-transcribed to cDNA using the PrimeScript RT Master Mix (Takara Bio, Kusatsu, Japan), and quantified using the THUNDERBIRD SYBR qPCR mix (Toyobo, Osaka, Japan). Then, qPCR was performed using StepOne Plus Real-Time PCR (Applied Biosystems, Carlsbad, CA, USA), with the following primers: Glyceraldehyde-3-phosphate dehydrogenase (forward, 5′-CATGGCCTTCCGTGTTCCTA-3′; reverse, 5′-GATGCCTGCTTCACCACCTT-3′) and *TNF-*α (forward, 5′-ACGTGGAACTGGCAGAAGAG-3′; reverse, 5′-TGAGGGTCTGGGCCATAGAA-3′). Threshold cycle values of each gene were normalized to those of glyceraldehyde-3-phosphate dehydrogenase. The fold change relative to the control group was calculated using the ΔΔCT method.^(30)^

### Enzyme-linked immunosorbent assay (ELISA)

Mice were anesthetized with a mixture of medetomidine (0.75 mg/kg, i.p.), midazolam (4 mg/kg, i.p.), and butorphanol (5 mg/kg, i.p.). Blood samples were collected, and the mice were euthanized by cervical dislocation. These samples were immediately collected 4 h after saline or LPS administration or RS exposure. Blood samples were allowed to clot and centrifuged to obtain serum. Each striatum was minced using surgical scissors in the ice-cold radioimmunoprecipitation assay buffer (FUJIFILM Wako Pure Chemical, Osaka, Japan) containing 1% phenylmethanesulfonyl fluoride. The suspension was sonicated on ice using CELL DISRUPTOR 200 (Branson, Danbury, CT, USA) and centrifuged at 16,000 × *g* for 10 min at 4 °C. After centrifugation, protein concentration in the supernatant was measured, and it was stored at –80 °C until ELISA. TNF-α concentration was calculated using the TNF-α ELISA kit for mice (Quantikine ELISA; R&D Systems, Minneapolis, MN, USA), according to the manufacturer’s instructions. Samples yielding TNF-α concentrations below the assay detection limit could not be reliably quantified and were therefore excluded from quantitative group comparisons.

### Immunohistochemistry

Mice were euthanized by cervical dislocation and brain samples for immunostaining were obtained 4 h after saline or LPS administration or RS exposure. The mice were transcardially perfused with PBS, followed by perfusion with 4% paraformaldehyde in PB (0.1 M; pH 7.2) under anesthesia with a mixture of medetomidine (0.75 mg/kg, i.p.), midazolam (4 mg/kg, i.p.), and butorphanol (5 mg/kg, i.p.). Their brains were dissected and placed in the same fixative overnight at 4 °C and cryoprotected in 30% sucrose/0.1 M PB solution. The brains were frozen and cut into 40-µm-thick sections using a freezing microtome (Leica SM2000R, Wetzlar, Germany). To prevent deformation, the sections were free-floated and blocked with 0.1% Triton X-100 and 3% bovine serum albumin for 30 min at 4 °C. After blocking, the sections were incubated with the rabbit polyclonal anti-ionized calcium-binding adaptor molecule 1 (Iba-1) antibody (1:500; Wako, Osaka, Japan) at 4 °C for 1 d. After washing with PBS, the sections were incubated with the Alexa555-labeled donkey anti-rabbit IgG antibody (1:300; Life Technologies, Carlsbad, CA, USA) for 30 min at room temperature. Then, the sections were counterstained with 4’,6-diamidino-2-phenylindole and mounted with the ProLong Diamond Antifade Mountant (Invitrogen, Thermo Fisher Scientific, Carlsbad, CA, USA). Fluorescent images were observed under the Olympus BX50 Fluorescence Microscope (Olympus, Tokyo, Japan) and captured using a camera and imaging software. For each mouse, five representative images (40× objective lens and 10× eyepiece) were acquired, sampling from across the striatum. Subsequently, the number of microglial cells and their cell body area were analyzed as previously described,^(31)^ with minor modifications. Analysis was conducted using the ImageJ Software (NIH Image, Bethesda, MD, USA). A threshold for positive staining was determined for each image including all cell bodies and processes, while excluding background staining. The results are reported as the average percentage area with the positive threshold in all representative images.

### Isolation of microglia

Mice were euthanized by cervical dislocation and whole-brains of four mice were extracted, pooled, and minced 4 h after saline or LPS administration or RS exposure. The cell suspension was prepared using a neural tissue dissociation kit (Miltenyi Biotec, Bergisch Gladbach, Germany), according to the manufacturer’s protocol. Then, the cell suspension was incubated with CD11b microbeads (Miltenyi Biotec, Bergisch Gladbach, Germany) in a custom buffer (prepared by dissolving 1.86 g of ethylenediaminetetraacetic acid in 1 L of PBS and adding 50 mL of fetal bovine serum) for 20 min at 4 °C. The cells were washed with the custom buffer, resuspended, and transferred to an LS column (Miltenyi Biotec, Bergisch Gladbach, Germany) under a magnetic field. Positively selected (CD11b+ microglia) cells were collected, and total RNA was extracted. RNA samples with concentrations below 5 ng/µL were excluded prior to reverse transcription. Subsequently, RT-qPCR was performed.

### Data and Statistical analyses

All data were analysed by experimenters blinded to the experimental conditions. Experimental groups were designed to have approximately equal sizes. For behavioural tests, the sample size was at least 5 per group. All graphs, calculations, and statistical analyses were conducted using the GraphPad Prism software version 8.0 for Mac (GraphPad Software, San Diego, CA, USA) and JMP Pro 17.0.0 (SAS Institute Inc., Cary, NC, USA). Data are represented as the mean ± standard error of the mean. Data normality was assessed using the Shapiro–Wilk test. No statistical outliers were removed from the datasets. Statistical analyses were performed using one-way or two-way analysis of variance (ANOVA), depending on the experimental design. Two-way ANOVA was used when two independent variables were included within the same experimental design, and their interaction was of interest. For repeated-measures data, two-way repeated-measures ANOVA was used. One-way ANOVA was used for experiments involving a single independent factor. When significant effects were detected, appropriate post hoc tests (Dunnett’s or Tukey’s multiple comparisons test) were applied. Dunnett’s test was used when comparisons were made against a single control group, whereas Tukey’s test was used for all other multiple comparisons. Statistical significance was set at p < 0.05.

## Results

### Effects of LPS and RS on behavioural sensitization

First, we investigated the effects of LPS and RS on behavioural sensitization. Control and METH-sensitized mice groups were created by administering saline and METH (1 mg/kg) for five consecutive days (Fig. 1A). One week after the fifth METH dose, METH-induced locomotor activity was measured as the primary behavioural outcome (Fig. 1A). In the repeated METH-treated group, a progressive enhancement of locomotor responses was observed over the course of administrations (Day 1, Day 5, and Day 12 post-withdrawal) (Fig. S2A–C). Notably, the locomotor activity on Day 12 was significantly higher than that on Day 1, demonstrating the establishment of behavioural sensitization (p < 0.01; Fig. S2B). To evaluate the behavioural changes in the presence of inflammation, saline or 1 μg/kg LPS administration or 2-h RS exposure was performed 4 h before the behavioural tests. Locomotor activity was significantly higher in the METH-sensitized mice compared to control mice in the saline condition confirming the expected enhancement of locomotor responses in METH-sensitized mice (Two-way ANOVA: F_METH_ [1,38] = 16.6 [p < 0.001], F_SAL, LPS, RS_ [2,38] = 4.65 [p < 0.05], and F_METH+SAL, LPS, RS_ [2,38] = 5.53 [p < 0.01]; Simple effects analysis under the SAL condition: F [1,38] = 24.7 [p < 0.001]; Fig. 1B). LPS and RS did not affect the METH-induced locomotor activity in control mice (p > 0.05; Fig. 1B, left). Both LPS and RS significantly suppressed METH-induced locomotion in the METH-sensitized mice (p < 0.05; Fig. 1B, right). The time-course of locomotor activity in the METH-sensitized mice, analysed in 5 min bins, is shown in Fig. 1C.

**Fig. 1.**
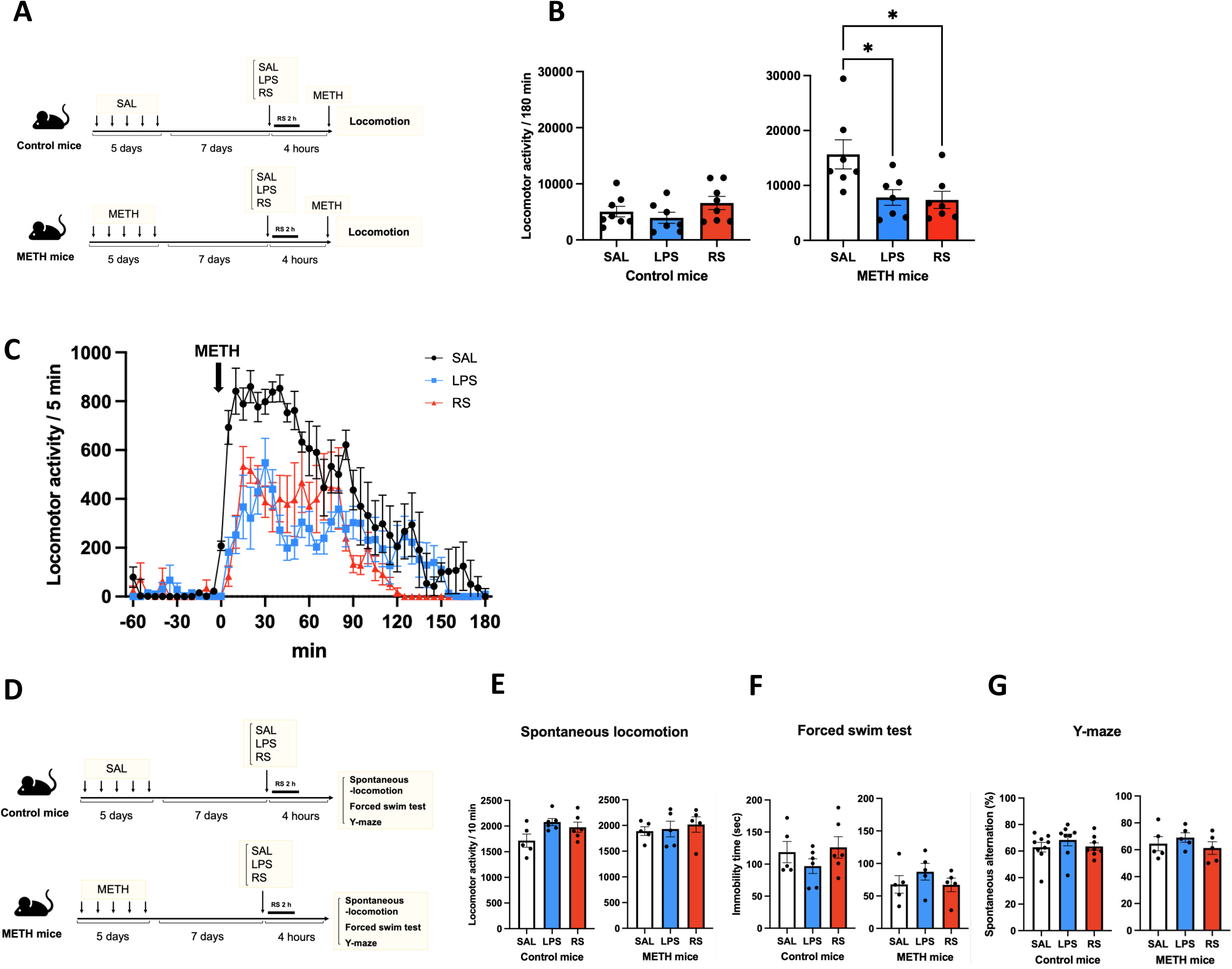
Effects of lipopolysaccharides (LPS) and restraint stress (RS) on behavioural sensitization. **(A)** Schematic timeline of the behavioural sensitization test. **(B)** Effects of LPS and RS on the behavioural sensitization of control (n = 8 [SAL], 7 [LPS], and 8 [RS]; left) and METH-sensitized (n = 7; right) mice. **(C)** Time-course of locomotor activity measured in 5-min bins of METH-sensitized mice (n = 7). **(D)** Schematic timeline of the spontaneous locomotion, forced swim, and Y-maze tests. **(E)** Effects of LPS and RS on the spontaneous locomotion of control (n = 5 [saline], 6 [LPS], and 6 [RS]; left) and METH-sensitized (n = 5; right) mice. **(F)** Effects of LPS and RS on the forced swim test results of control (n = 5 [SAL], 6 [LPS], and 6 [RS]; left) and METH-sensitized (n = 5; right) mice. **(G)** Effects of LPS and RS on the Y-maze test results of control (n = 9 [SAL], 8 [LPS], and 8 [RS]; left) and METH-sensitized (n = 5; right) mice. Two-way analysis of variance (ANOVA) was used to examine the main effects of the model (Control, METH) and treatment (SAL, LPS, RS), as well as their interaction. When a significant interaction was found, simple effects analyses were conducted to assess the effect of the model within each treatment condition. Additionally, separate One-way ANOVAs were performed within each model to examine the effect of treatment condition, followed by Tukey’s post hoc tests when appropriate. Dots indicate the individual mice in all experimental groups. Data are represented as the mean ± standard error of the mean (SEM). Statistical significance was set at *p < 0.05. **(B)** Two-way ANOVA: F_METH_ [1,38] = 16.6 [p < 0.001], F_SAL, LPS, RS_ [2,38] = 4.65 [p < 0.05], and F_METH+SAL, LPS, RS_ [2,38] = 5.53 [p < 0.01]; Simple effects analysis under the SAL condition: F [1,38] = 24.7 [p < 0.001]; One-way ANOVA in the control mice: p > 0.05; Dunnett’s test in the METH-sensitized mice: SAL vs. LPS [p < 0.05], SAL vs. RS [p < 0.05]. **(E)** Two-way ANOVA: F_METH_ [1,26] = 0.69 [p > 0.05], F_SAL, LPS, RS_ [2,26] = 1.47 [p > 0.05], and F_METH+SAL, LPS, RS_ [2,26] = 0.27 [p > 0.05]; One-way ANOVA in the control mice: p > 0.05; One-way ANOVA in the METH-sensitized mice: p > 0.05. **(F)** Two-way ANOVA: F_METH_ [1,26] = 14.7 [p < 0.001], F_SAL, LPS, RS_ [2,26] = 0.08 [p > 0.05], and F_METH+SAL, LPS, RS_ [2,26] = 2.22 [p > 0.05]; One-way ANOVA in the control mice: p > 0.05; One-way ANOVA in the METH-sensitized mice: p > 0.05. **(G)** Two-way ANOVA: F_METH_ [1,33] < 0.01 [p > 0.05], F_SAL, LPS, RS_ [2,33] = 1.15 [p > 0.05], and F_METH+SAL, LPS, RS_ [2,33] = 0.14 [p > 0.05]; One-way ANOVA in the control mice: p > 0.05; One-way ANOVA in the METH-sensitized mice: p > 0.05. ns, not significant. SAL, saline; METH, methamphetamine.

Additionally, we evaluated spontaneous locomotion, immobility during the forced swim test, and alternation behaviour in the Y-maze test during the same experimental timeline using separate mouse cohorts for each test (Fig. 1D). No significant changes in spontaneous locomotor activity were observed across the three groups (saline, LPS, and RS) in both the control and METH-sensitized mice (Two-way ANOVA: F_METH_ [1,26] = 0.69 [p > 0.05], F_SAL, LPS, RS_ [2,26] = 1.47 [p > 0.05], and F_METH+SAL, LPS, RS_ [2,26] = 0.27 [p > 0.05]; One-way ANOVA in the control mice: p > 0.05; One-way ANOVA in the METH-sensitized mice: p > 0.05 ; Fig. 1E). In the forced swim test, immobility duration was significantly reduced in METH-sensitized mice compared to control mice, although there were no significant differences among SAL, LPS, and RS conditions in either the control or METH-sensitized mice (Two-way ANOVA: F_METH_ [1,26] = 14.7 [p < 0.001], F_SAL, LPS, RS_ [2,26] = 0.08 [p > 0.05], and F_METH+SAL, LPS, RS_ [2,26] = 2.22 [p > 0.05]; One-way ANOVA in the control mice: p > 0.05; One-way ANOVA in the METH-sensitized mice: p > 0.05 ; Fig. 1F). In the Y-maze test, spontaneous alternation percentage showed no significant differences between control mice and METH-sensitized mice, nor among SAL, LPS, and RS conditions in either group (Two-way ANOVA: F_METH_ [1,33] < 0.01 [p > 0.05], F_SAL, LPS, RS_ [2,33] = 1.15 [p > 0.05], and F_METH+SAL, LPS, RS_ [2,33] = 0.14 [p > 0.05]; One-way ANOVA in the control mice: p > 0.05; One-way ANOVA in the METH-sensitized mice: p > 0.05 ; Fig. 1G). Since LPS and RS significantly affected behavioural sensitization only in the METH-sensitized mice but not in the control mice, subsequent analyses focusing on the mechanisms underlying this effect were conducted in the METH-sensitized mice group.

### Roles of inflammation-related factor inhibitors in the LPS- and RS-induced suppression of behavioural sensitization

To examine the association between LPS- and RS-induced behavioural changes and inflammation-related factors, TLR4, TNF-α, and COX-2 inhibitor were administered immediately before LPS administration and RS exposure (Fig. 2A). When TAK-242, a TLR4 inhibitor, was combined with LPS, behavioural reduction observed with LPS pre-treatment was not detected in the LPS + TAK-242 group (Two-way ANOVA: F_LPS_ [1,17] = 15.0 [p < 0.01], F_TAK-242_ [1,17] = 14.4 [p < 0.01], and F_LPS+TAK-242_ [1,17] = 6.96 [p < 0.05]; Tukey’s test: LPS + VEH vs. LPS + TAK-242 [p < 0.05]; Fig. 2B, left). Similarly, combining TAK-242 with RS mitigated the decreased locomotor activity observed after RS pre-treatment (Two-way ANOVA: F_RS_ [1,20] = 9.94 [p < 0.01], F_TAK-242_ [1,20] = 26.9 [p < 0.001], and F_RS+TAK-242_ [1,20] = 16.1 [p < 0.001]; Tukey’s test: RS + VEH vs. RS + TAK-242 [p < 0.001]; Fig. 2B, right).

**Fig. 2.**
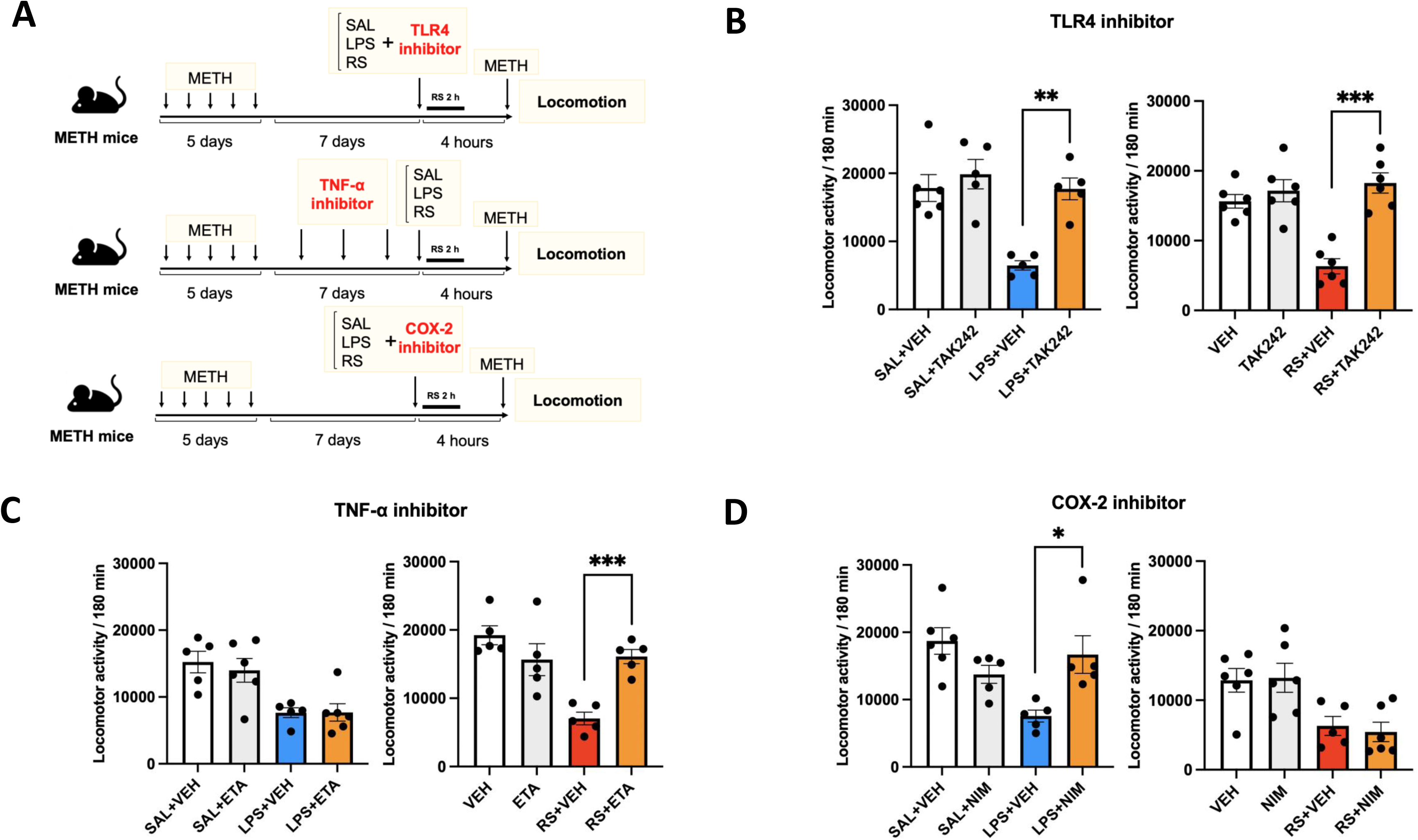
Roles of inflammation-related factor inhibitors in the LPS- and RS-induced suppression of behavioural sensitization. **(A)** Schematic timeline of the behavioural sensitization test. **(B)** Effects of TAK-242, a toll-like receptor 4 (TLR4) inhibitor, on METH-sensitized mice treated with LPS (n = 6 [SAL + VEH], 5 [SAL + TAK-242], 5 [LPS + VEH], and 5 [LPS + TAK-242]; left) and RS (n = 6; right). **(C)** Effects of etanercept (ETA), a tumour necrosis factor (TNF)-α inhibitor, on the METH-sensitized mice treated with LPS (n = 5 [SAL + VEH], 6 [SAL + ETA], 5 [LPS + VEH], and 6 [LPS + ETA]; left) and RS (n = 5; right). **(D)** Effects of cyclooxygenase-2 (COX-2) inhibitor, on the METH-sensitized mice treated with LPS (n = 6 [SAL + VEH], 5 [SAL + NIM], 5 [LPS + VEH], and 5 [LPS + NIM]; left) and RS (n = 6 [SAL + VEH], 6 [SAL + NIM], 5 [RS + VEH], and 6 [RS + NIM]; right). Dots indicate the individual mice in all experimental groups. Data are represented as the mean ± SEM. Statistical significance was set at *p < 0.05. **p < 0.01 and ***p < 0.001 vs. LPS and RS groups treated with and without TAK-242, ETA, or nimesulide (NIM). Two-way ANOVA, followed by Tukey’s test, was used for multiple-group comparisons. **(B)** Left: Two-way ANOVA: F_LPS_ [1,17] = 15.0 [p < 0.01], F_TAK-242_ [1,17] = 14.4 [p < 0.01], and F_LPS+TAK-242_ [1,17] = 6.96 [p < 0.05]; Tukey’s test: LPS + VEH vs. LPS + TAK242 [p < 0.05]. Right: Two-way ANOVA: F_RS_ [1,20] = 9.94 [p < 0.01], F_TAK-242_ [1,20] = 26.9 [p < 0.001], and F_RS+TAK-242_ [1,20] = 16.1 [p < 0.001]; Tukey’s test: RS + VEH vs. RS + TAK-242 [p < 0.001]. **(C)** Left: Two-way ANOVA: F_LPS_ [1,18] = 19.6 [p < 0.001], F_ETA_[1,18] = 0.70 [p > 0.05], and F_LPS+ETA_ [1,18] < 0.01 [p > 0.05]. Right: Two-way ANOVA: F_RS_ [1,16] = 14.7 [p < 0.01], F_ETA_ [1,16] = 3.21 [p > 0.05], and F_RS+ETA_ [1,16] = 17.0 [p < 0.0001]; Tukey’s test: RS + VEH vs. RS + ETA [p < 0.01]. **(D)** Left: Two-way ANOVA: F_LPS_ [1,17] = 4.58 [p < 0.05], F_NIM_ [1,17] = 1.18 [p > 0.05], and F_LPS+NIM_ [1,17] = 13.5 [p < 0.01]; Tukey’s test: LPS + VEH vs. LPS + NIM [p < 0.05]. Right: Two-way ANOVA: F_RS_ [1,19] = 17.8 [p < 0.01], F_NIM_ [1,19] = 0.02 [p > 0.05], and F_RS+NIM_ [1,19] = 0.13 [p > 0.05]. ETA, etanercept; NIM, nimesulide; SAL, saline; METH, methamphetamine; VEH, vehicle.

When TNF-α inhibitor etanercept (ETA) was administered prior to LPS treatment, locomotor activity in the LPS + ETA group was significantly lower than that in the control group (Two-way ANOVA: F_LPS_ [1,18] = 19.6 [p < 0.001], F_ETA_ [1,18] = 0.70 [p > 0.05], and F_LPS+ETA_ [1,18] < 0.01 [p > 0.05]; Fig. 2C, left). In contrast, etanercept combined with RS prevented the RS-induced suppression of locomotor activity. Locomotor activity in the RS + ETA group was significantly higher than that in the RS + vehicle group but comparable to that in the vehicle group (Two-way ANOVA: F_RS_ [1,16] = 14.7 [p < 0.01], F_ETA_ [1,16] = 3.21 [p > 0.05], and F_RS+ETA_ [1,16] = 17.0 [p < 0.0001]; Tukey’s test: RS + VEH vs. RS + ETA [p < 0.01]; Fig. 2C, right).

When COX-2 inhibitor nimesulide (NIM) was administered prior to LPS treatment, decrease in locomotor activity observed in the LPS + vehicle group was not noted in the LPS + NIM group (Two-way ANOVA: F_LPS_ [1,17] = 4.58 [p < 0.05], F_NIM_ [1,17] = 1.18 [p > 0.05], and F_LPS+NIM_ [1,17] = 13.5 [p < 0.01]; Tukey’s test: LPS + VEH vs. LPS + NIM [p < 0.05]; Fig. 2D, left). However, locomotor activity in the LPS + NIM group was comparable to that in the saline + vehicle group, suggesting that nimesulide prevents the LPS-induced suppression of locomotion. In contrast, when nimesulide was administered prior to RS treatment, decrease in locomotor activity observed in the RS + vehicle group remained unchanged in the RS + NIM group (Two-way ANOVA: F_RS_ [1,19] = 17.8 [p < 0.01], F_NIM_ [1,19] = 0.02 [p > 0.05], and F_RS+NIM_ [1,19] = 0.13 [p > 0.05]; Fig. 2D, right). Notably, locomotor activity in the RS + NIM group was significantly lower than that in the vehicle group (p < 0.05) but comparable to that in the RS + vehicle group (p > 0.05), suggesting that COX-2 inhibition does not affect the RS-induced suppression of locomotion.

### Effects of LPS and RS on METH-induced striatal dopamine levels

We investigated whether LPS and RS affect the striatal extracellular dopamine levels in response to METH administration. Compared to the saline group, LPS group showed no significant difference in the extracellular dopamine levels after METH administration (p > 0.05; Fig. 3A, left). Notably, dopamine levels in the LPS group peaked at similar levels and followed a time course comparable to that in the saline group, with no significant differences detected at any measured time point (p > 0.05; Fig. 3A, left). Conversely, the RS group showed significantly lower extracellular dopamine levels than the saline group at 40, 80, and 120 minutes after METH administration (p < 0.05; Fig. 3A, left). Moreover, area under the curve (AUC) for changes in dopamine levels over 3 h following METH administration showed no significant difference between the LPS and saline groups (p > 0.05; Fig. 3A, right). In contrast, the AUC of the RS group was significantly lower than that of the saline group (p < 0.01; Fig. 3A, right). Furthermore, administration of ETA as TNF-α inhibitor prior to RS pre-treatment significantly inhibited the RS-induced suppression of METH-induced increase in extracellular dopamine levels (p < 0.05; Fig. 3B, left). Dopamine levels in the RS + ETA group were significantly higher than those in the RS group (p < 0.05; Fig. 3B, left), reaching levels comparable to those in the saline group (p > 0.05; Fig. 3B, left). AUC for changes in dopamine levels in the RS + ETA group was significantly higher than that in the RS group (p < 0.05; Fig. 3B, right) but comparable to that in the saline group (p > 0.05; Fig. 3B, right).

**Fig. 3.**
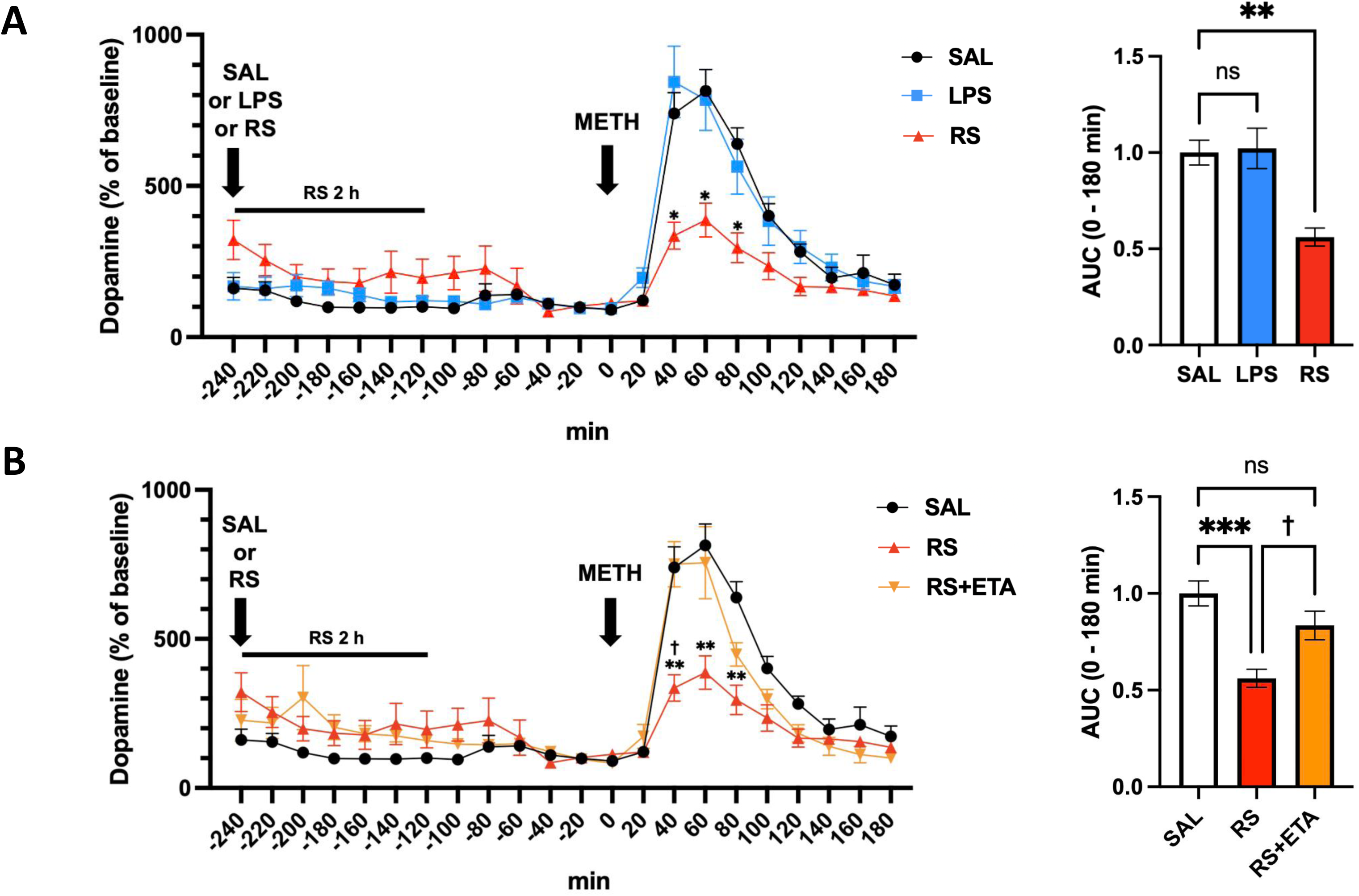
Effects of LPS and RS on methamphetamine (METH)-induced striatal dopamine responses. **(A)** Left: Effects of LPS and RS on METH-induced striatal dopamine responses (n = 5 [SAL], 6 [LPS], and 5 [RS]). Right: Area under the curve (AUC) for changes in dopamine levels over 3 h following METH administration. Two-way repeated-measures ANOVA followed by post hoc multiple comparisons was used for time-course data (0-180 min), and one-way ANOVA followed by Dunnett’s test was used for AUC analysis. Two-way RM ANOVA: F _time_ [1.613, 20.97] = 64.32, p < 0.001; F _SAL, LPS, RS_ [2,23] = 4.445, p = 0.0338; F _time × SAL, LPS, RS_ [18,117] = 5.075, p < 0.001. **(B)** Left: Effects of RS and RS + ETA on METH-induced striatal dopamine responses (n = 5). Right: AUC for changes in dopamine levels over 3 h following METH administration. Two-way repeated-measures ANOVA followed by post hoc multiple comparisons was used for time-course data (0-180 min), and one-way ANOVA followed by Tukey’s test was used for AUC analysis. Two-way RM ANOVA: F _time_ [2.409, 28.9] = 62.82, p < 0.001; F _SAL, RS, RS+ETA_ [2,12] = 6.740, p = 0.009; F _time × SAL, RS, RS+ETA_ [18,108] = 5.303, p < 0.001. Data are represented as the mean ± SEM. Statistical significance was set at *p < 0.05, **p < 0.01, and ***p < 0.001 RS vs SAL; **†**p < 0.05 RS vs RS+ETA. AUC, area under curve; ETA, etanercept; SAL, saline.

### Effects of LPS and RS on TNF-**α** expression levels

We examined the TNF-α expression levels in peripheral blood and the striatum. In peripheral blood, TNF-α protein levels were measured across all groups to assess the systemic inflammatory responses. TNF-α protein levels were elevated only in the LPS group, whereas its levels were below the detection limit in the other groups (Fig. 4A). In the striatum, TNF-α protein levels were significantly elevated in the RS group compared to those in the saline group (p < 0.05; Fig. 4B). Notably, TNF-α protein levels in the LPS group were not significantly different from those in the saline group.

**Fig. 4.**
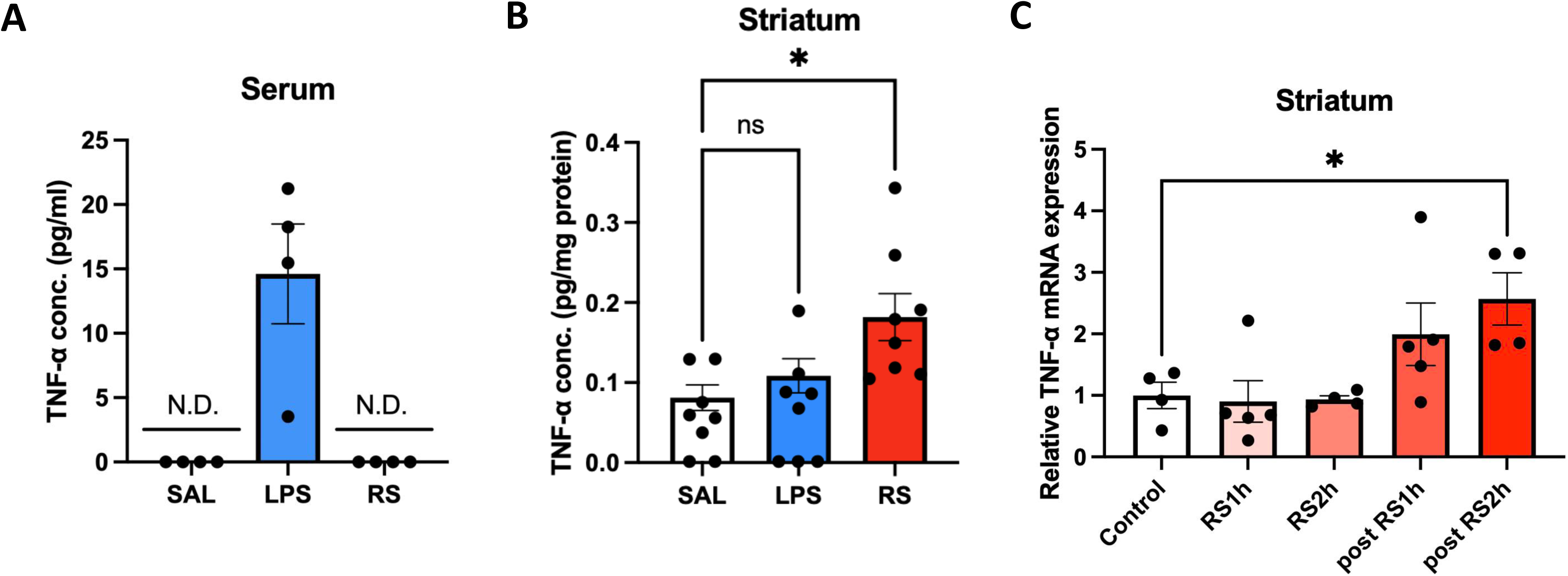
**Effects of LPS and RS on TNF-**α **expression levels. (A)** Effects of LPS and RS on TNF-α protein levels in peripheral blood (n = 4). **(B)** Effects of LPS and RS on TNF-α protein levels in the striatum (n = 8). **(C)** *TNF-*α mRNA expression levels in the striatum following RS exposure at various time points (n = 4 [control], 5 [RS1h], 4 [RS2h], 5 [post-RS1h], and 4 [post-RS2h]). One-way ANOVA, followed by Dunnett’s test, was used for multiple-group comparisons. **(B)** Samples with TNF-α levels below the detection limit were excluded from quantitative analysis (2 samples in SAL and 3 samples in LPS groups). Dots indicate the individual mice in all experimental groups. Data are represented as the mean ± SEM. Statistical significance was set at *p < 0.05 (**p < 0.01 and ***p < 0.001). N.D., not detected; SAL, saline.

Subsequently, *TNF-*α mRNA levels in the striatum were evaluated. We assessed the changes in *TNF-*α mRNA levels following RS exposure over time. *TNF-*α mRNA levels were increased by RS, showing a significant elevation 4 h after the initiation of RS exposure (post-RS2h) compared to those in the untreated group (p < 0.05; Fig. 4C).

### Effects of LPS and RS on striatal microglia

We hypothesized that the source of TNF-α production was microglia. To verify this, we evaluated the effects of LPS and RS on microglia. Fluorescence microscopic evaluation of Iba1, a microglial marker, revealed no significant changes in the Iba1-positive area of the striatum in response to LPS and RS treatment (p > 0.05; Fig. 5A and B). Similarly, number of Iba1-positive cells in the striatum was not significantly altered by either treatment (p > 0.05; Fig. 5C). To specifically investigate TNF-α production in microglia, the microglia were isolated from mouse whole-brains using magnetic bead-labelled anti-CD11b antibodies. *TNF-*α mRNA levels were subsequently measured in the isolated microglia. Notably, *TNF-*α mRNA levels were significantly increased in the RS group compared to those in the saline group (p < 0.05; Fig. 5D). However, *TNF-*α mRNA levels in the LPS group were not significantly different from those in the saline group (p > 0.05; Fig. 5D).

**Fig. 5.**
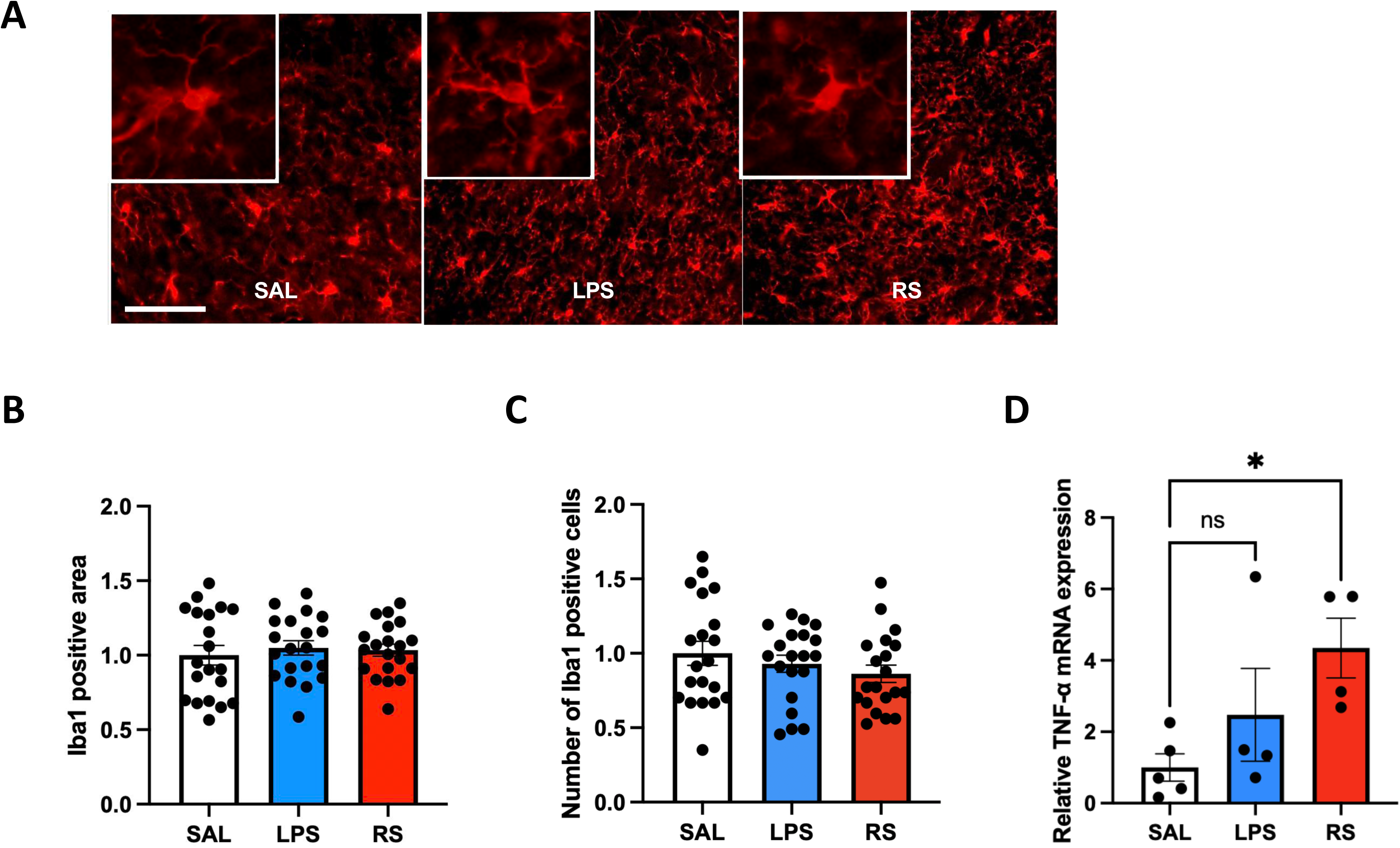
Effects of LPS and RS on microglia. **(A)** Representative immunostaining image for Iba1. Scale bars, 100 μm (low-power photos) and 10 μm (high-power photos). **(B)** Iba1-positive area in the striatum (n = 20). **(C)** Number of Iba1-positive cells in the striatum (n = 20). **(D)** Effects of LPS and RS on mRNA expression levels in separated microglia (N = 5 [SAL], 4 [LPS], and 4 [RS]). One-way ANOVA, followed by Dunnett’s test, was used for multiple-group comparisons. In Fig. 5B and C, each plot represents a single analysed image. As shown in Fig. 5D, data of four mice were calculated as a single value, represented by N = 1 in each plot. Data are represented as the mean ± SEM. Statistical significance was set at * p < 0.05 (**p < 0.01 and ***p < 0.001). ns, not significant.

## Discussion

In this study, LPS- or RS-induced acute inflammation induced under relatively mild, clinically relevant conditions suppressed behavioural sensitization in the METH-sensitized mice, without inducing other behavioural effects. Furthermore, despite both LPS and RS suppressing behavioural sensitization in the METH-sensitized mice, their underlying mechanisms were distinct: LPS exerted its effects via COX-2 without altering the striatal dopamine levels, whereas the effects of RS were dependent on microglia-derived TNF-α and associated with a significant reduction in the striatal dopamine levels.

In this study, acute inflammation suppressed behavioural sensitization in the METH-sensitized mice. Notably, this improvement was observed under extremely low-intensity inflammation, such as that induced by LPS (1 μg/kg) and a single 2-h RS session, which was far lower than the doses typically used in the inflammation models in animal studies.^(32)^ These findings suggest that even low-intensity inflammation improves behavioural sensitization in the METH-sensitized mice. To verify this, we conducted spontaneous locomotion, forced swim, and Y-maze tests, as high-intensity inflammation influences these behaviours.^(17, 33, 34)^ No significant changes were observed in these tests, confirming that the observed suppression of behavioural sensitization was not due to general motor impairments, depressive-like behaviours, or cognitive alterations. These findings suggest that the observed effects are unlikely to reflect a sickness behaviour-like state associated with high-intensity inflammation.

In this study, both LPS and RS attenuated behavioural sensitization in the METH-sensitized mice but did not affect the METH-induced hyperactivity in the control mice. METH-induced sensitization is widely used as a model of psychostimulant-induced psychosis and is relevant to certain aspects of schizophrenia-related pathophysiology. It is thought to involve dopaminergic mechanisms, including reverse tolerance of dopamine receptors.^(11, 14)^ This suggests that LPS and RS affect behavioural sensitization by modulating dopamine activity via inflammation. To verify this, we examined the inflammation-related factors involved in LPS- and RS-induced behavioural changes and assessed their impacts on the dopaminergic system.

Both LPS and RS attenuated behavioural sensitization via TLR4 activation. TLR4 is a key receptor involved in the immune responses to bacterial endotoxins, such as LPS, that initiates immune responses.^(35)^ It is considered to be involved in stress-induced neuroinflammation.^(35–37)^ Consistently, TLR4 was involved in both LPS-induced immune responses and RS-induced stress-related neuroimmune-mediated behavioural changes in the METH-sensitized mice in this study.

We also examined the downstream effectors of TLR4. Notably, RS-induced suppression of behavioural sensitization was associated with increased TNF-α levels and decreased dopamine levels in the striatum. TNF-α plays a key role in dopaminergic regulation,^(38)^ suggesting a mechanistic link between RS-induced neuroinflammation and dopamine modulation. Additionally, TNF-α promotes the uptake of extracellular dopamine via TNF receptor 1 and dopamine transporter,^(38, 39)^ thereby reducing the extracellular dopamine availability. Moreover, administration of TNF-α in the METH-sensitized model reduces the striatal extracellular dopamine levels and behavioural sensitization.^(38)^ These findings suggest that RS attenuates behavioural sensitization by directly modulating the striatal dopamine levels via TNF-α signalling.

In contrast to that induced by RS, suppression of behavioural sensitization induced by LPS occurred without significant changes in striatal TNF-α and dopamine levels. In this study, LPS-induced behavioural suppression was blocked by a COX-2 inhibitor, indicating that LPS effects were mediated via COX-2 activation.

Prostaglandins enhance the gamma-aminobutyric acid (GABA)ergic input into the substantia nigra via E-type prostanoid receptors on GABAergic neurons, thereby indirectly modulating their dopaminergic activity.^(40, 41)^ In this study, COX-2 activation possibly contributed to this process by promoting prostaglandin signalling in the METH-sensitized mice. These results suggest that RS and LPS suppress behavioural sensitization via different mechanisms, with RS acting via TNF-α-mediated dopamine modulation and LPS exerting its effects via the COX-2 pathway.

Next, to determine whether microglia are involved in the mechanisms by which acute inflammation improves behavioural sensitization, we focused on TNF-α, which was directly linked to dopaminergic modulation. Stress can affect microglial function in ways that likely vary with the duration and intensity of exposure, with acute stress lasting hours to less than a day and chronic stress lasting days to weeks thought to engage partially distinct neuroimmune responses.^(42)^ Acute stress has been reported to induce rapid microglial responses, including increased production of inflammatory cytokines, changes in inflammatory markers on the surface, and enhanced responsiveness to subsequent stimuli.^(42)^ In the present study, we observed a significant increase in *TNF-*α mRNA levels in the microglia isolated from RS-treated mice, without any morphological changes. Our findings suggest that acute RS induces functional changes in microglia without increasing their cell number or cell body area, consistent with previous reports on single-session short-term stress.^(21)^ In contrast, LPS elevated the TNF-α levels in peripheral blood, but not in the striatum and microglia. As LPS and inflammatory cytokines do not cross the blood–brain barrier,^(43)^ their effects on the brain are possibly mediated via peripheral immune signalling. Relatively low-dose LPS (10 μg/kg) activates the COX-1–prostaglandin pathway in endothelial cells, thereby promoting prostaglandin E2 production and depressive-like behaviours.^(44)^ This study observed COX-2 activation instead of COX-1 activation, suggesting that prostaglandins derived from endothelial cells contribute to LPS-induced neuroinflammation.

Collectively, these findings suggest that RS and LPS suppress behavioural sensitization in the METH-sensitized mice via distinct mechanisms: RS via direct microglial effects and LPS via peripheral immune signalling without altering microglial states. This highlights the diverse cellular pathways involved in inflammation-induced behavioural modulation. Our findings provide insights into the mechanisms by which acute inflammation improves behavioural sensitization.

In this study, acute inflammation effectively suppressed behavioural sensitization in the METH-sensitized mice. Importantly, these findings were obtained under relatively mild and clinically relevant inflammatory conditions, suggesting that even low-intensity inflammatory responses can modulate behavioural sensitization. These findings provide new insights into how inflammatory responses may influence psychosis-related outcomes, with potential relevance to positive symptoms of schizophrenia, although validation in other models and symptom-related behaviour will be necessary. Furthermore, the distinct mechanisms identified for LPS and RS raise the possibility that different inflammatory pathways may differentially influence dopaminergic function and behavioural outcomes. Although we focused on TNF-α and COX-2 based on their established roles in neuroimmune–dopamine interactions, other molecular mechanisms may also contribute to the observed effects. Future studies using broader molecular approaches, such as high-throughput transcriptomic analyses, will be important to clarify the full range of pathways engaged by distinct inflammatory triggers.

In conclusion, acute inflammation induced under relatively mild conditions suppressed behavioural sensitization in METH-sensitized mice, with distinct mechanisms depending on the inflammatory trigger. Although further studies are required to establish clinical relevance, our findings provide a conceptual framework for understanding the context-dependent roles of inflammation in psychosis-related conditions including schizophrenia and may contribute to the development of novel therapeutic strategies targeting inflammation-related pathways.

## Data Availability Statement

Raw data supporting the findings of this study are available upon request from the corresponding author.

## Acknowledgements

We thank the researchers who contributed to this study, as well as the staff of the Animal Research Facility and the Department of Psychiatry, Graduate School of Medicine, Hokkaido University, for their valuable support. We would also like to thank Editage (http://www.editage.jp) for English language editing.

## Author Contributions

RS, SI, and KI conceived the study and designed the experiments. RS, SI, KI, RM, MT, and SM performed the experiments. MO and MK advised on the microglial analysis procedure. RS and SI analyzed the data and prepared the figures. RS and SI wrote and edited the manuscript with input from all authors. NH, IK, and TK supervised the study.

