## Supplemental Figure for "Acute inflammation-mediated attenuation of behavioural sensitization in methamphetamine-sensitized mice via distinct COX-2 and TNF-α pathways"

### SUPPLEMENTAL INFORMATION

### Supplementary Figure

**Fig. S1**

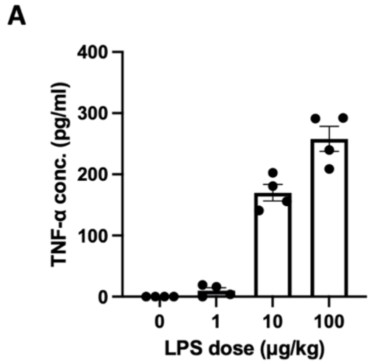


**Fig. S2**

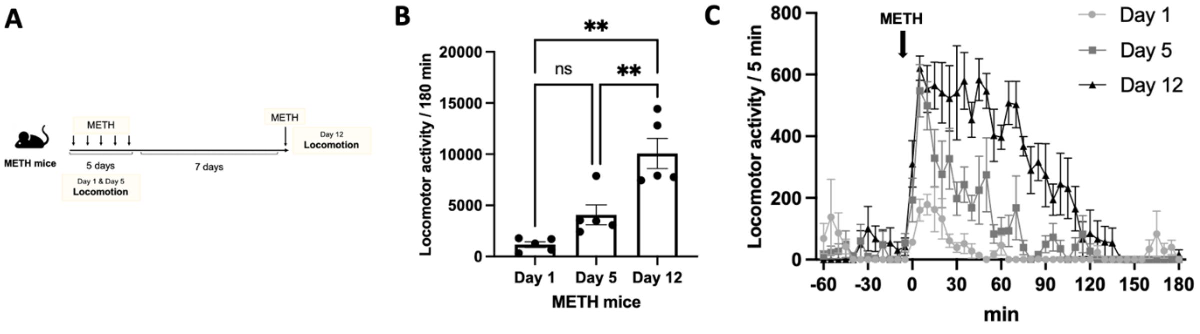

### Supplementary Figure Legends

**Fig. S1. Effects of various lipopolysaccharide (LPS) doses on the tumour necrosis factor (TNF)-α protein levels in peripheral blood. (A)** Effects of various LPS doses on TNF-α protein levels in peripheral blood (n = 4). Dots indicate the individual mice in all experimental groups. Data are represented as the mean ± standard error of the mean (SEM).

**Fig. S2. Effects of repeated METH administration on METH induced locomotor activity. (A)** Schematic timeline of METH induced locomotion test. **(B)** METH induced locomotor activity on Day 1, Day 5, and Day 12 (n = 5). **(C)** Time-course of locomotor activity measured in 5-min bins (n = 5). Dots indicate the individual mice in all experimental groups. Data are represented as the mean ± standard error of the mean (SEM). One-way ANOVA, followed by Tukey's test, was used for multiple-group comparisons. Statistical significance was set at $*p < 0.05$, $**p < 0.01$. ns, not significant. METH, methamphetamine.

### Supplementary Methods

#### LPS dose selection

The LPS dose was determined based on preliminary experiments (Fig. S1A) and selected as the minimal dose that produced detectable peripheral immune activation. TNF-α was used as a representative inflammatory cytokine based on previous reports of its elevation in patient with schizophrenia and its involvement in dopaminergic regulation as demonstrated in experimental studies.[(1), (2)] Because baseline cytokine levels differ between species, direct quantitative comparison between mice and humans is limited; therefore, to avoid excessive inflammation, cytokine responses were interpreted in a relative framework, aligning the magnitude of differences between inflammatory and control conditions in the experimental model with those observed between patients and healthy controls in clinical studies.[(3)] In dose–response experiments (Supplementary Fig. 1A), administration of 1 μg/kg of LPS resulted in detectable peripheral TNF-α elevation, whereas higher doses (≥10 μg/kg) induced substantially greater cytokine responses, demonstrating a dose-dependent effect. Higher doses of LPS are known to induce excessive inflammatory responses and sickness-like behaviour,[(4), (5)] whereas the lowest dose tested (1 μg/kg) was sufficient to produce detectable TNF-α elevation. Therefore, 1 μg/kg was selected as the effective dose to model mild immune activation while avoiding confounding effects associated with higher doses.
